# Resting-state aperiodic neural activity as a novel objective marker of daytime somnolence

**DOI:** 10.1101/2022.08.07.503118

**Authors:** Zachariah R. Cross, Teng Yuan Kang, Kurt Lushington, Paroma Sarkar, Parmjit Singh, Sonya Johnston, Aeneas Yeo, Alex Chatburn

**Affiliations:** Cognitive Neuroscience Laboratory - Australian Research Centre for Interactive and Virtual Environments, University of South Australia, Adelaide, Australia; Sleep & Respiratory Failure Program, Department of Thoracic Medicine, Royal Adelaide Hospital, Central Adelaide Local Health Network, Adelaide, Australia

**Keywords:** excessive daytime sleepiness, EEG, aperiodic slope, slow oscillation-spindle coupling, multiple sleep latency test

## Abstract

**Study Objectives:** Current assessment of excessive daytime somnolence (EDS) requires subjective measurements such as the Epworth Sleepiness Scale (ESS), and/or resource intensive sleep laboratory investigations. Recent work ^1,2^ has called for more non-performance-based measures of EDS. One promising non-performance-based measure of EDS is the aperiodic component of electroencephalography (EEG). Aperiodic (non-oscillatory) activity reflects excitation/inhibition ratios of neural populations and is altered in various states of consciousness, and thus may be a potential biomarker of hypersomnolence.

**Methods:** We retrospectively analysed EEG data from patients who underwent a Multiple Sleep Latency Test (MSLT) and determined whether aperiodic neural activity is predictive of EDS. Participants having undergone laboratory polysomnogram and next day MSLT were grouped into MSLT+ (*n* = 26) and MSLT– (*n* = 33) groups (mean sleep latency of < 8min and > 10min, respectively) and compared against a non-clinical (Control) group of participants (*n* = 26).

**Results:** While the MSLT+ and MSLT– groups did not differ in their aperiodic activity, the Control group had a significantly flatter slope and larger offset compared to both MSLT+ and MSLT– groups. Logistic regression machine learning predicted group status (i.e., symptomatic, non-symptomatic) with 90% accuracy based on the aperiodic slope while controlling for age. Slow oscillation-spindle coupling was also significantly stronger in the Control group relative to MSLT+ and MSLT– groups.

**Conclusions:** Our results provide first evidence that aperiodic neural dynamics and sleep-based cross-frequency coupling is predictive of EDS, thereby providing a novel avenue for basic and applied research in the study of sleepiness.

## 1. Introduction

It is broadly accepted that sleep must serve a vital, biological purpose, or be evolution’s greatest error ^3^. So too sleepiness, or somnolence, defined as both the subjective experience of accumulated sleep need and the likelihood of falling asleep at any given moment, must be a marker of an important biological process in much the same way that pain is the subjective experience of nociception. Excessive daytime sleepiness (EDS), or hypersomnolence, comes with inherent risk to both the individual, such as poorer mental health outcomes, and the community, including greater risk of motor vehicle accidents and occupational errors and injuries ^4,5^. Despite the health risks associated with hypersomnolence, the direct measurement of neurobiological markers of sleepiness are difficult to obtain ^6,7^. For example, current gold standards rely on subjective questionnaires, such as the Epworth Sleepiness Scale (ESS), and the Multiple Sleep Latency Test (MSLT), both of which are not without flaws (e.g., ^8^), with concerns regarding the expense, access and reliability of the MSLT ^9^. Methodological issues with current measures of EDS were highlighted in a recent review ^1^ and editorial ^2^ in the flagship journal *SLEEP*, both of which called upon the need for non-subjective and performance-based measures of EDS. Here, we present data which indicates that somnolence can be read directly from the human electroencephalogram (EEG) via the aperiodic component.

Somnolence is explained in the two-process model ^10,11^ as resulting from the interaction of homeostatic and circadian drives to sleep. That is, sleep pressure is the accumulation of sleep need that increases across wake and dissipates across sleep ^12^, and is influenced by circadian timing ^10^. Mechanistically, the accumulation of sleep need is proposed to result from long-term potentiation (LTP) due to incidental learning and information processing during wakeful behaviour, resulting in saturation of synaptic strength in the cerebral cortex ^13–15^. Data supporting this position are drawn from studies showing that: i) neural firing rates increase as a function of time spent awake, and then decrease over sleep ^16^; ii) molecular and electrophysiological markers of LTP are positively related to hours of wakefulness, while markers of global synaptic depression are linked with time spent in subsequent sleep ^14^; and, iii) using transcranial magnetic stimulation to directly measure neural firing responses, there is a noted increase in synaptic strength in both excitatory and inhibitory cortical connections in response to prolonged wakefulness ^17^. Taken together, this body of work demonstrates that somnolence may result from plastic changes in the brain that accumulate over wakefulness. Therefore, a direct measure of brain activity should contain information related to the need for sleep in the individual.

Research indicates several potential markers of somnolence in the EEG ^7^. Proposed oscillatory markers of somnolence include alpha attenuation ^18^, theta/alpha ratio ^19^ and changes in theta power ^12,20^. Theta power, in particular, increases in magnitude as a function of time spent awake and correlate with slow wave activity in the first subsequent sleep cycle^12^ in line with model predictions ^10,11,13^.

Despite this body of research, neural measures of somnolence have not been widely adopted, and current approaches to the measurement of somnolence involve either full-day behavioural testing, or subjective survey instruments ^6,7^. There are several potential reasons for this, including difficulties in interpreting individual differences in the EEG and lack of normal data around this ^7^, a lack of understanding as to what oscillatory techniques here may actually be measuring (i.e., a spectral increase may reflect a compensatory mechanism in response to somnolence, not somnolence *per se* ^6,21^, and noted circadian influences in the oscillatory markers which would need to be accounted for unless they confound the signal ^22–24^.

One potential novel approach for the direct measurement of somnolence is through the aperiodic component of the EEG. This approach has several advantages, including that aperiodic activity is intuitively linked to neurobiologically-informed models of sleep need as it demarks modulations in neuronal excitability ^25^, and has been shown to reliably differ between both healthy and sleep disordered patients ^26^ as well as between wakeful, anaesthetised, REM and NREM brain states ^25^.

Given the utility of aperiodic activity in distinguishing between states of consciousness and between health and disease, we aim to describe aperiodic factors in the EEG as markers of somnolence. We calculated the aperiodic slope and intercept from resting state EEG data from patients diagnosed with excessive daytime somnolence via MSLT designated MSLT+ for this cohort, patients who presented with excessive daytime somnolence but who did not meet diagnostic criteria on the MSLT (MSLT–), and healthy individuals who did not report excessive somnolence (Control). Using mixed-effects regression models, as well as machine learning approaches and advanced measures of sleep micro and macro-state variables, we present a thorough investigation of the utility of aperiodic EEG markers in characterising daytime somnolence, and potential links to resulting functional impairment.

## 2. Method

### 2.1. Participants and design

To obtain data from a symptomatic cohort we reviewed patients who were referred to the Royal Adelaide Hospital (RAH) Sleep Disorders Laboratory Sleep & Respiratory Failure Service for an overnight in-laboratory polysomnography (PSG) and next day MSLT between August 2014 and August 2017. Patients were included if they completed an overnight PSG and proceeded to next day MSLT. Participants were required to have ≥ 2 min (4 epochs) of resting awake EEG data in both PSG and all MSLT naps to enable reliable extraction of the aperiodic component.

This cohort of patients was split into the objectively sleepy group (MSLT+) which was defined as patients with a mean sleep latency (MSL) of <8 min, and patients without demonstrable objective sleepiness who had an MSL of >10min (MSLT–; ^27,28^). Patients with an “intermediate” MSL between 8 and 10min were not included to further stratify the two groups. Additionally, patients were excluded from analysis if they did not have sufficient quality EEG data of at least two minutes in a resting awake state. Other exclusion criteria were major psychiatric co-morbidity apart from depression, significant neurologic disability (including acquired or traumatic brain injury), use of typical or atypical anti-psychotic medications, prescribed stimulants or wakefulness promoting agents (e.g., Dexamfetamine, Methylphenidate, Modafinil and/or Armodafinil), a positive urinary drug screen, or untreated obstructive sleep apnoea (defined as an AHI > 10/hr).

Data were collected on baseline demographics (Table 1), use of medications that may potentially affect sleep stage, ESS, and PSG and MSLT parameters which were all completed at the time of study as per the sleep laboratory protocol (Figure 1A). The MSLT was performed as per the AASM recommendations ^28^. Four to five nap opportunities were provided during daylight hours with central and occipital electroencephalogram (EEG), chin muscle electromyogram (EMG), bilateral electro-oculogram (EOG), and electrocardiogram (ECG) activity recorded with calibrations performed at the beginning of each nap opportunity. The use of these data was approved by the Central Adelaide Local Health Network Human Research (CALHN ID: 15132) and University of South Australia’s Human Ethics (204266; 204778) Committees.

**Table 1.** Demographic and clinical variables of study groups.

| Clinical Variables | Controls<br>(n = 26) | MSLT –<br>(n = 33) | MSLT +<br>(n = 26) |
| --- | --- | --- | --- |
| Age (years) | 42 (22.80) | 38.80 (16.30) | 45.5 (18.00) |
| Gender (% female) | 73.08 | 69.70 | 69.23 |
| ESS | 5.33 (4.06) | 10.40 (4.60) | 12.40 (5.61) |

**Figure 1.**
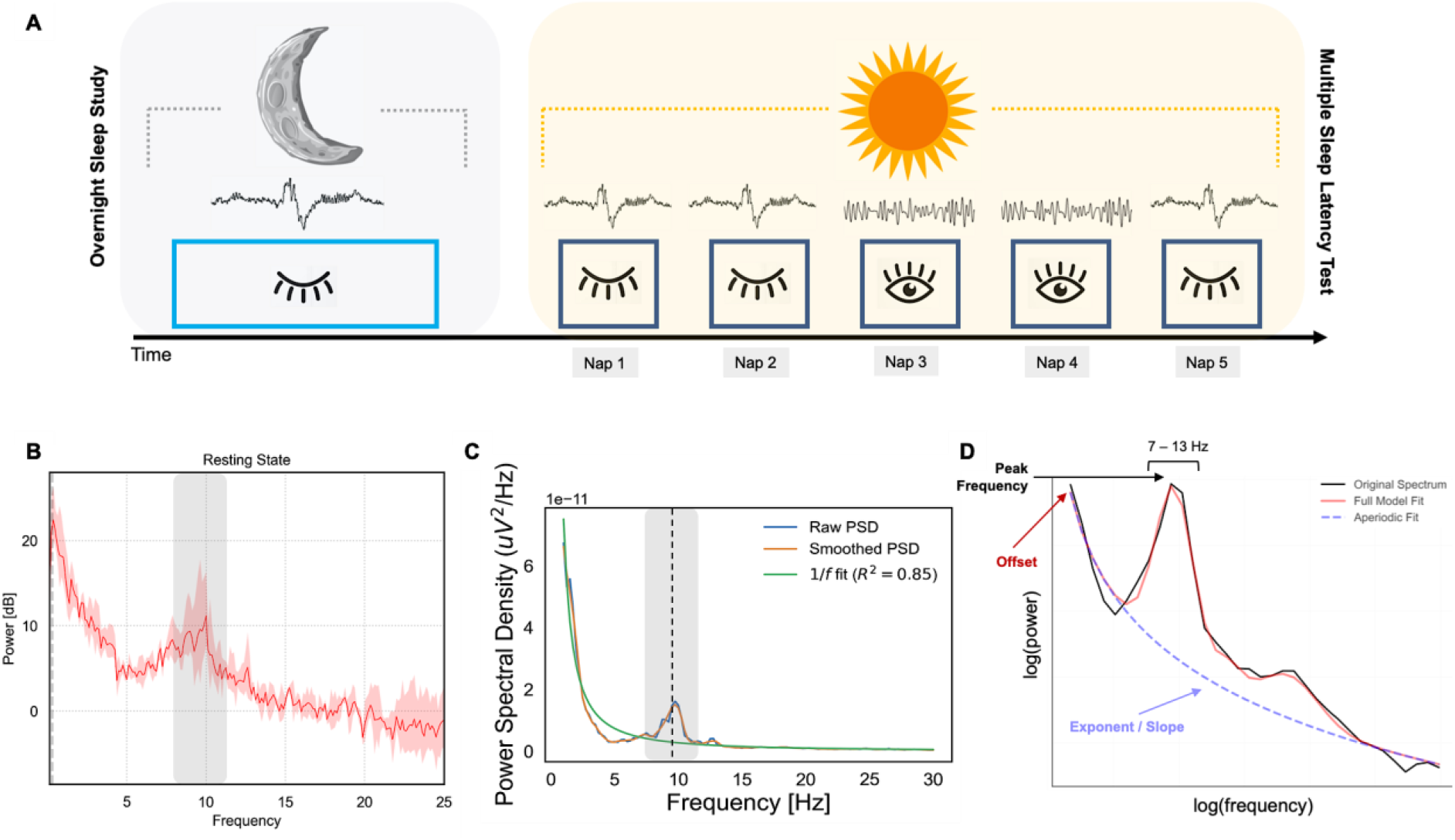
Schematic of Multiple Sleep Latency Test and resting-state-derived EEG metrics. (A) Subjects undertook five nap bouts throughout a day, following a monitored night of polysomnography (left; light blue rectangle). Open and closed eyes indicate if sleep was obtained on a nap bout (dark blue squares).

A group of healthy participants who had no reported sleep problems were used as a control group in analyses examining the aperiodic component using resting daytime EEG. Participants in the Control group were aged 18 – 80 years, monolingual native English speakers, were not visually impaired and did not report any medical disorders (Table 1). Control group participants also reported not taking any sleep-related medication, not having taken recreational drugs in the last six months, and no current diagnoses of neurological or psychological conditions, including past or current diagnosis of traumatic brain injury.

A second group of healthy participants (*n* = 17; M_age_ = 23.60; 8 females) were used for analyses examining sleep-related EEG. This group was based on archival data from ^29^. Participants were aged 18 – 39 years and met the same inclusion criteria described above for the first control group.

### 2.2. EEG Pre-Processing

#### 2.2.1. Resting-state EEG

Patient data (MSLT–, MSLT+) were retrospectively analysed from archival PSG and MSLT data from the Royal Adelaide Hospital Sleep Disorders Laboratory between August 2014 and 2017. EEG recordings were obtained from attended polysomnography (PSG) and MSLT using Profusion PSG 4 software (E-series, Compumedics, Abbortsford, Vic Australia). The Control group data were obtained from either a LiveAmp or actiCHamp system (Brain Products GmbH, Gilching, Germany) with 32 or 64 active Ag/AgCl electrodes mounted in an elastic cap (actiCap or BrainCap, Brain Products GmbH, Gilching, Germany). Electrode placement for both patient (MSLT+, MSLT–) and control (aperiodic and sleep-based healthy comparisons) groups followed the 10/20 system. Horizontal and vertical eye movements and blinks were monitored with electrodes placed above and slightly below the left eye. EEG channels from the Control group were amplified using a BrainVision amplifier (LiveAmp 32 or actiCHamp base unit 5001, Brain Products, GmbH) at either a 500 Hz or 1000 Hz sampling rate. All EEG pre-processing and analysis was performed using MNE-Python (Gramfort et al., 2013). Raw data were band-pass-filtered from 0.1 to 30 Hz (zero-phase, hamming windowed finite impulse response). Data were then re-referenced to the average of left and right mastoid electrodes and resampled to 250 Hz to match the sampling rate of the patient recordings.

#### 2.2.2. Sleep EEG

Sleep data were scored by expert sleep scientists according to standardised criteria ^30^ using Compumedics Profusion 4 software (Abbortsford, Vic, Australia). The EEG was viewed with a high-pass filter of 0.3 Hz and a low-pass filter of 35 Hz. The following sleep parameters were calculated: total sleep time, sleep onset latency, wake after sleep onset, time (minutes) and percent of time spent in each sleep stage (N1, N2, N3 and R). Slow oscillation-spindle (SO-spindle) coupling strength were extracted via the YASA toolbox ^31^ implemented in MNE-Python ^32^ based on previously used methods (e.g., ^33,34^).

For pre-processing of the sleep EEG, the EEG signal was re-referenced to linked mastoids and filtered from 0.1 – 30 Hz using a digital phase-true FIR band-pass filter. Data were then epoched into 30 second bins and subjected to a multivariate covariance-based artifact rejection procedure ^35,36^.

### 2.3. EEG Data Analysis

#### 2.3.1. Neural aperiodic exponent

To estimate the 1/*f* power-law exponent from resting-state EEG recordings, we used the irregular-resampling auto-spectral analysis method (IRASA v1.0; ^37^) implemented in the YASA toolbox ^31^ in MNE-Python. The (negative) exponent summarising the slope of aperiodic spectral activity was calculated by fitting a linear regression to the estimated fractal component in log-log space. For a full mathematical description of IRASA, see ^37^ (also see ^38^).

#### 2.3.2. NREM slow oscillations and spindle coupling

For slow oscillations, continuous NREM EEG data were filtered using a digital phase-true FIR band-pass filter from 0.3 – 2 Hz with a 0.2 Hz transition band to detect zero crossing events that were between 0.3 – 1.5 s in length, and that met a 75 to 500 microvolt criterion. These artifact-free epochs were then extracted from the raw EEG signal.

We also estimated SO-spindle coupling activity (for a detailed description of this method, see ^33^). We first filtered the normalized SO trough-locked data into the SO component (0.1 – 1.25 Hz) and extracted the instantaneous phase angle after applying a Hilbert transform. Then we filtered the same trials between 12 – 16 Hz and extracted the instantaneous amplitude from the Hilbert transform. For every participant and epoch, we detected the maximal sleep spindle amplitude and corresponding SO phase angle at electrode C3. The resultant vector length (mean vector; coupling strength) across all NREM events were then determined using circular statistics implemented in the pingouin package ^39^. The Rayleigh test was used to test for circular non-uniformity with *p* < 0.01.

## 3. Statistical Analysis

Data were imported into *R* version 4.0.2 (R Core Team, 2020) and analysed using linear mixed-effects regressions fit with restricted maximum likelihood (REML) estimates using *lme4* ^40^. For models predicting differences in the aperiodic slope and offset, and in ESS scores, Condition (Control, MSLT– and MSLT+) was specified as a fixed effect, and Sex (Male, Female) and Age were treated as covariates. Similarly, for models examining differences in sleep-related EEG (e.g., SO density, SO-spindle coupling), Condition was specified as a fixed effect, while Age was treated as a covariate. For all EEG-based models, participant ID and electrode were modelled as random effects on the intercept to account for between-subject and topographic differences in aperiodic estimates, respectively. Type II Wald χ2-tests from the *car* package ^41^ were used to provide *p*-value estimates, while effects were plotted using the package *effects* ^42^ and *ggplot2* ^43^. Outliers were isolated using Tukey’s method, which identifies outliers as exceeding ± 1.5 × inter-quartile range. Categorical factors were sum-to-zero contrast coded, such that factor level estimates were compared to the grand-mean ^44^. Further, an 83% confidence interval (CI) threshold was used given that this approach corresponds to the 5% significance level with non-overlapping estimates ^45,46^. In the visualisation of effects, non-overlapping CIs indicate a significant difference at *p* < 0.05.

## 4. Results

### 4.1. Resting-state aperiodic neural activity predicts clinical status

In modelling the aperiodic slope, we revealed a significant main effect of Group, with the Control group having a flatter 1/*f* slope compared to both the MSLT– (*β* = -0.19, *se* = 0.06, *p* = 0.001) and MSLT+ (*β* = -0.14, *se* = 0.06, *p* = 0.02; Figure 2A) groups. Critically, the MSLT– and MSLT+ groups did not differ in their 1/*f* slope estimates (*β* = -0.05, *se* = 0.04, *p* = 0.15). A similar pattern emerged when modelling differences in the aperiodic offset: while there was no difference in the offset between the MSLT+ and MSLT– groups (*β* = 0.17, *se* = 0.16, *p* = 0.28), the Control group had a larger offset than the MSLT+ (*β* = 0.82, *se* = 0.24, *p* = 0.001; Figure 2B) and MSLT– (*β* = 0.99, *se* = 0.24, *p* < 0.001) groups.

**Figure 2.**
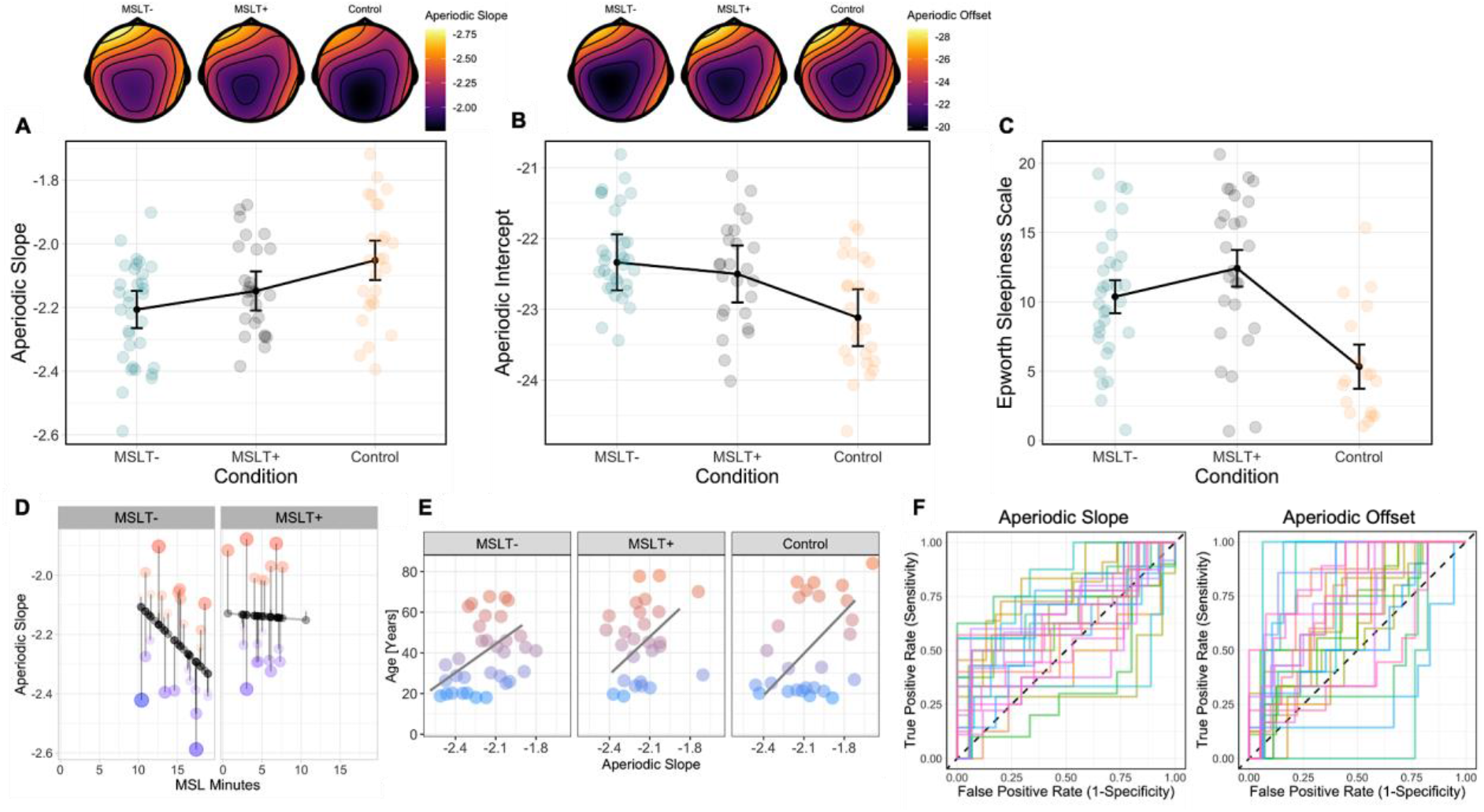
Differences in aperiodic activity and self-perceived sleepiness between study groups. (A) Differences in the aperiodic slope (y-axis; higher values denote a flatter slope) between the MSLT–, MSLT+ and Control groups (x-axis). (B) Represents the same as (A) but for the aperiodic offset. Topographical distribution of the aperiodic slope and offset between study groups are given above (A) and (B), respectively. (C) Differences in self-perceived sleepiness quantified by the ESS (y-axis; higher values denote more self-perceived sleepiness) between the MSLT–, MSLT+ and Control groups (x-axis). For A – C, bars represent the 83% confidence interval around group-level expected marginal mean estimates. Dots represent individual data points per subject for aggregated data. (D) Relationship between the aperiodic slope (y-axis) and MSL in minutes (x-axis; higher values denote a longer time to fall asleep). Points along the line of best fit indicate predicted values, while points off the line represent residual data, with larger points indicating further distance from predicted values. (E) Relationship between age (in years; y-axis; higher values denote older age) and the aperiodic slope (x-axis) across each study group. (F) Region over the curve plot representing the logistic machine learning model for the aperiodic slope (left) and offset/intercept (right). Each line represents one of the 25 bootstrapped resamples of the training data.

For analyses regarding self-perceived sleepiness (quantified as ESS), we found a main effect of Group (Figure 2C): while there was no difference between the MSLT– and MSLT+ groups (*β* = 2.05, *se* = 1.28, *p* = 0.11), the Control group had significantly lower ESS scores (*β* = -5.04, *se* = 1.43, *p* < 0.001). However, when adding the aperiodic slope into the model, there was no significant main effect of Group (*β* = 19.03, *se* = 17.77, *p* = 0.29), Slope (*β* = -2.21, *se* = 5.55, *p* = 0.70), or interaction between Group and Slope (*β* = 11.72, *se* = 8.41, *p* = 0.17) on self-perceived sleepiness.

Based on these observations, we performed machine learning analyses to determine whether we could predict clinical status based on resting-state derived aperiodic measures. Given that the MSLT+ and MSLT– groups did not differ in either the 1/*f* slope and offset, we collapsed these groups together, resulting in a two-level factor of Symptomatic (MSLT+, MSLT–) and non-symptomatic (Control). Data were separated into a training and test set, retaining 75% of the data for training and 25% for testing. We then created bootstrapped resamples of the training data to evaluate the logistic regression models, which predicted Group (Control, MSLT+, MSLT–) from 1/*f* Slope and Age, and the 1/*f* Offset and Age. The model containing the 1/*f* slope performed well on the test data with an accuracy estimate of 0.90 and region under the curve estimate of 0.90, while the 1/*f* offset model yielded an accuracy and region under the curve estimate of 0.70 and 0.66, respectively. From this, we were able to predict with 90% accuracy whether an individual is symptomatic or non-symptomatic based on the combination of the 1/*f* slope and age. For a visualisation of both machine learning models, see Figure 2F.

### 4.2. SO-spindle coupling differs between individuals with and without EDS

We examined whether there were differences in sleep-related oscillatory dynamics between those with MSLT+, MSLT–, and Controls (i.e., no diagnosed sleep disorders). We focussed on three cardinal measures of slow oscillations, namely slow oscillation (SO) density, SO peak-to-peak amplitude, and the slope of SOs. We also quantified SO-spindle coupling, which has shown to change across the lifespan ^33^, as well as differ between those with and without psychiatric disorders ^47^. While controlling for age, there was no significant difference in SO density (χ^2^(2) = 0.94, *p* = 0.62), peak-to-peak amplitude (χ^2^(2) = 0.71, *p* = 0.70), or SO slope (χ^2^(2) = 2.25, *p* = 0.32) between the three groups. There was also no significant difference in SO-spindle coupling density (*F*(2,146) = 0.22, *p* = 0.80; for visualisation of these metrics across groups, see Figure 4). Note that a mixed-effects regression did not converge for SO-spindle density, and thus a simple linear regression was performed (i.e., without the random intercept of subject). However, there was a significant difference in SO-spindle coupling strength (χ^2^(2) = 287.45, *p* < 0.001) between the three groups, which is resolved in Figure 4E. While the MSLT+ and MSLT– groups did not differ in SO-spindle coupling strength (*β* = 0.0002, *se* = 0.002, *p* = 0.93), the Control group had stronger SO-spindle coupling strength than the MSLT+ (*β* = 0.06, *se* = 0.003, *p* < 0.001) and MSLT– (*β* = 0.06, *se* = 0.004, *p* < 0.001) groups. There was also a similar effect of Coupling Phase, whereby there was a significant difference between groups in the preferred spindle amplitude to SO phase (χ^2^(2) = 41.57, *p* < 0.001). Here, maximal spindle amplitude occurred after the peak of the SO in the Control group, while spindle amplitude was maximal prior to the SO peak for both the MSLT– and MSLT+ groups (Figure 4F). For a summary of sleep parameter metrics between groups, see Table 2, while for a visualisation of SO-spindle coupling activity, see Figure 3.

**Table 2.** Sleep parameters of study groups.

| Sleep Measures | Controls | Patients | | $F(p, \eta_p^2)$ |
| --- | --- | --- | --- | --- |
| | ($n = 17$) | MSLT- ($n = 33$) | MSLT+ ($n = 26$) | |
| TST | 400.00 (67.02) | 389.00 (61.70) | 405.00 (59.7) | 1.37 (.24, .02) |
| SOL | 15.23 (12.23) | 29.80 (23.2) | 25.30 (21.8) | 0.30 (.60, .01) |
| WASO | 52.64 (55.60) | 58.10 (47.00) | 44.80 (40.60) | 1.51 (.22, .02) |
| N1 | 10.05 (8.21) | 4.43 (3.85) | 4.00 (2.80) | 0.11 (.73, .004) |
| N2 | 49.52 (10.36) | 51.10 (11.2) | 55.70 (8.80) | 2.93 (.09, .05) |
| N3 | 25.84 (9.60) | 28.00 (12.5) | 22.00 (8.85) | 3.66 (.06, .06) |
| REM | 14.57 (8.56) | 16.50 (7.10) | 17.80 (6.57) | 0.48 (.49, .008) |
| SO Density | 12.90 (7.16) | 7.85 (7.53) | 6.92 (6.70) | 0.45 (.64, .20) |
| SO Peak-to-Peak | 119.64 (18.24) | 108.45 (19.91) | 109.54 (24.80) | 0.37 (.69, 0.09) |
| SO Slope | 477.41 (117.54) | 393.41 (114.73) | 397.60 (108.13) | 1.17 (.31, .20) |
| Coupling Density | 9.26 (7.49) | 6.60 (7.37) | 5.65 (6.70) | 2.90 (.06, .04) |
| Coupling Strength | 0.23 (0.01) | 0.17 (0.01) | 0.17 (0.00) | **345.92 (<.001, 0.82)** |
| Coupling Phase | -0.27 (0.34) | 0.26 (0.41) | 0.17 (0.38) | **21.65 (<.001, 0.23)** |

*Note.* Values presented in parentheses are standard deviation. TST = total sleep time; SOL = sleep onset latency; WASO = wake after sleep onset; N1 = stage 1; N2 = stage 2; N3 = slow wave sleep; REM = rapid eye movement sleep. Values for each sleep stage represent the percent of time spent in each stage. $\eta_p^2$ = partial eta squared.

**Figure 3.**
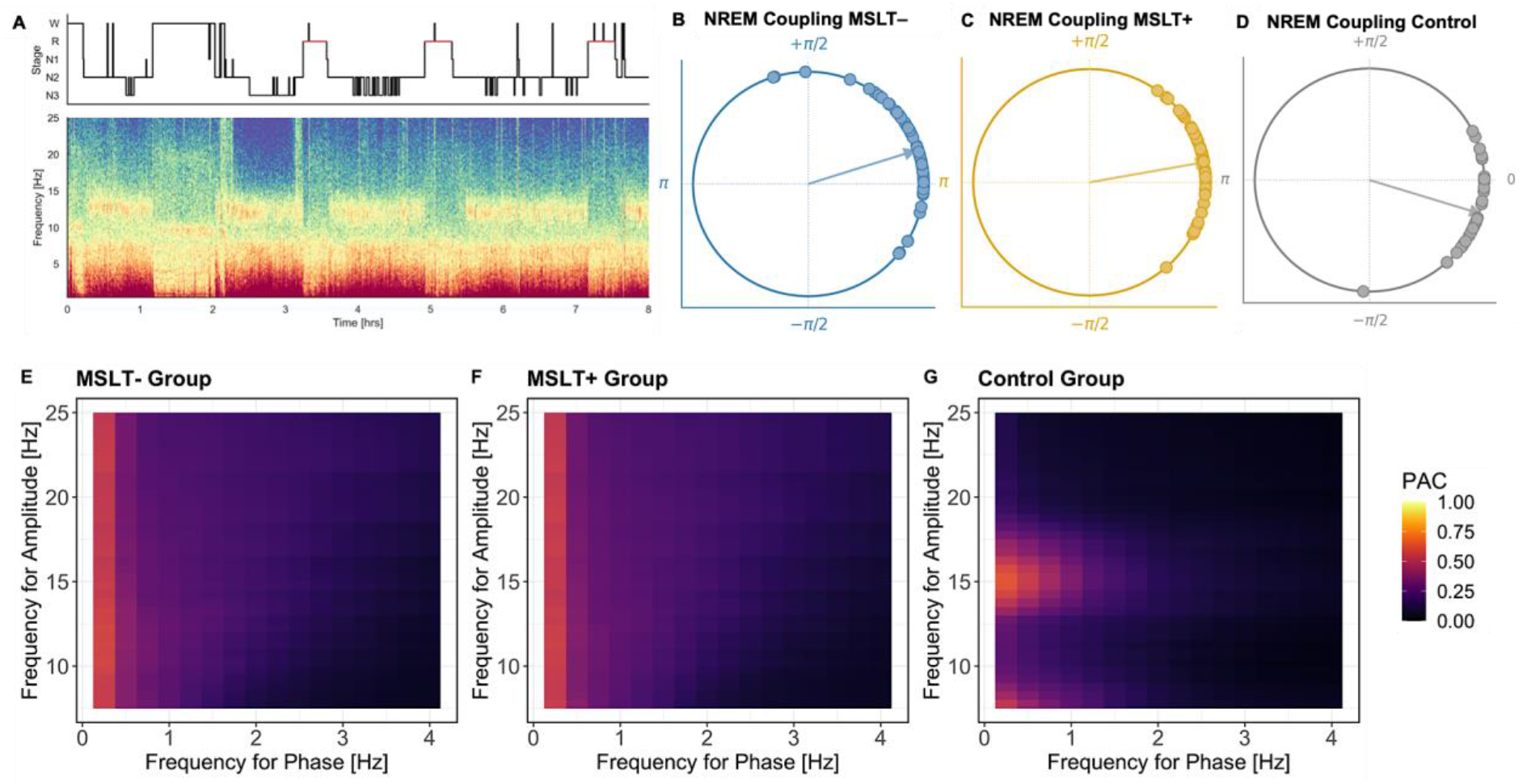
Sleep neurophysiology and NREM SO-spindle coupling activity between study groups. (A) Hypnogram and full-night multi-taper spectrogram for a single participant from channel Cz. (B – D) Preferred phase of SO-spindle coupling for each study group. Circles indicate individual participant estimates. Note that 0 represents the peak of the SO. (E – G) Group-level comodulagram illustrating the frequency for phase (x-axis) and frequency for power (y-axis) during NREM sleep SO-spindle epochs for each study group. PAC = phase amplitude coupling.

**Figure 4.**
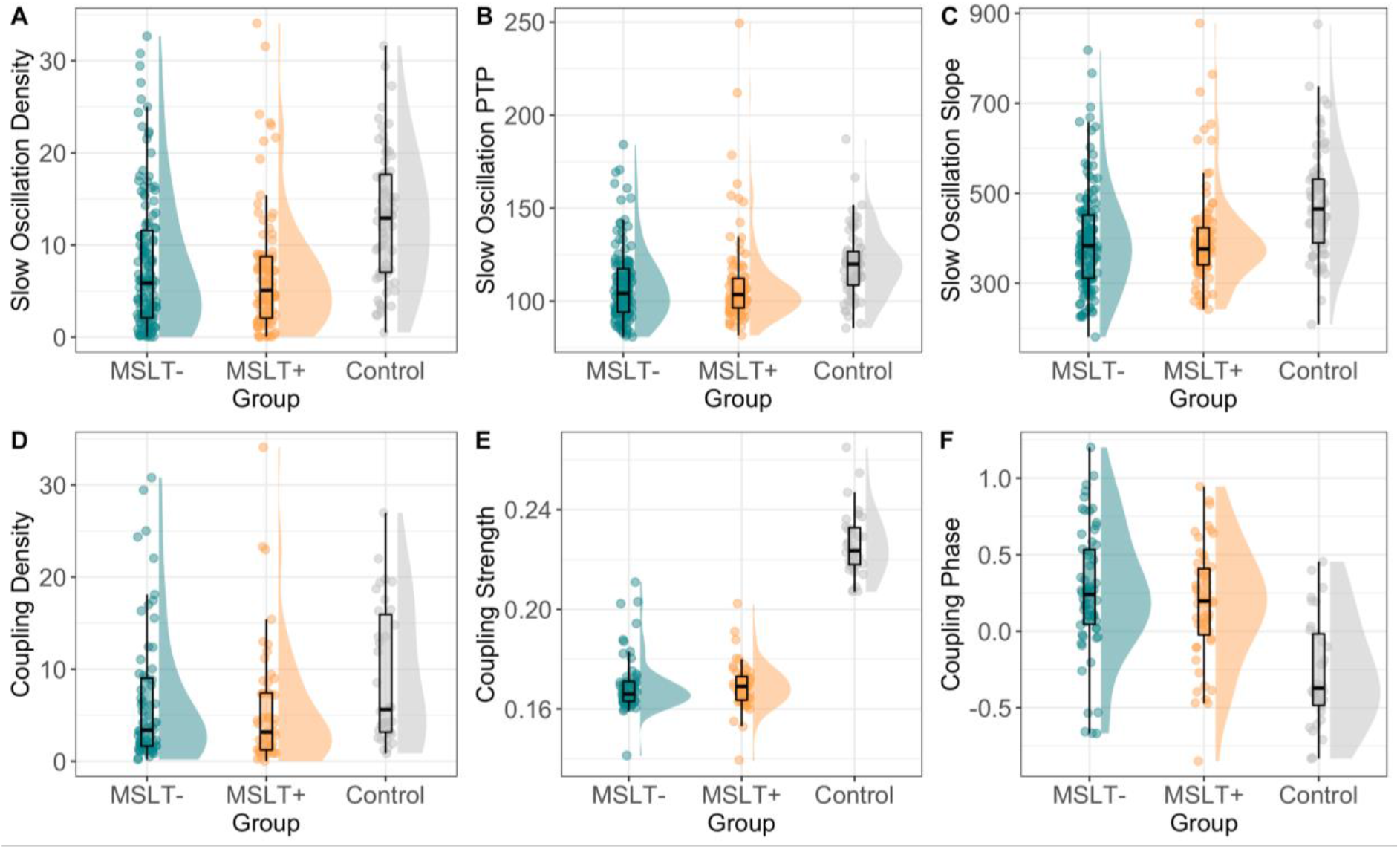
Sleep microstructural activity between study groups. For each plot, study group (MSLT–, MSLT+ and Control) is represented on the x-axis, while sleep microstructural activity is shown on the y-axis. In each plot, thick horizontal lines indicate the median; lower and upper hinges correspond to the first and third quartiles, respectively; lower and upper whiskers extend to the furthest estimate within 1.5 × interquartile range from the lower and upper hinges, respectively.

## 5. Discussion

In the present study we aimed to determine whether resting-state and sleep-related EEG metrics differentiate between individuals with and without excessive daytime somnolence (EDS). The aperiodic slope (a proxy for neural excitation/inhibition; ^48^) and slow-oscillation spindle (SO-spindle) coupling (related to neuronal transmission and subcortical-cortical connectivity; ^49^) were statistically predictive of EDS. Critically, these neural metrics were stronger predicters of EDS than the ESS and MSLT, offering a potentially novel objective measure for sleep physicians and scientists in the diagnosis and treatment of chronic sleepiness in patient populations. These findings also inform current understandings of the neurobiological basis of sleepiness, a phenomenon that serves a vital homeostatic function in both health and disease.

The slope of the aperiodic component has been recently described as a marker of homeostatic sleep need ^50^, based on theoretical accounts of neural activity leading to sleep need ^51,52^ and experimental results in animals and humans which have delineated processes of synaptic regulation as a function of sleep ^53–55^. Broadly, as the individual progresses through wakefulness, incidental information processing necessarily occurs, and this leads to saturation of synaptic connections in cortex. On a neural level, this leads to increased levels of excitability as a function of hours of wakefulness ^51,55,56^. Aperiodic metrics offer a unique opportunity to measure such processes non-invasively and cheaply, as they have been linked with the activity of excitatory and inhibitory neurons ^25,48,57^, with information in the aperiodic signal related to increases in both excitation and inhibition in the neural system. Therefore, to the extent that the mammalian drive to sleep is related to the accumulation of synaptic strength as a result of incidental information processing ^51,52^ leading to increased excitation ^16,53^ and that aperiodic measures reflect the E/I balance of the neural system ^25,58^, aperiodic measures may constitute a useful marker of currently experienced sleepiness in the individual, linking to homeostatic sleep need.

Our results broadly extend previous literature by demonstrating that the aperiodic slope can be used to classify cases of excessive daytime somnolence from healthy controls. These results are informative in terms of finding relatively cheap and efficient diagnostic markers of sleep disorders but are also in terms of understanding precisely what is measured by aperiodic markers in terms of sleep need. Sleep research typically differentiates both circadian and homeostatic sleep drives, and between sleep propensity and subjective sleepiness ^11,23,59,60^, and it is presently unclear how these linked drives can be read directly from the EEG, despite the brain being deeply impacted by them ^61,62^. Here, we note that the aperiodic slope is predictive of time to sleep onset in the MSLT (Figure 2D). As such, it presently appears that the information contained in the aperiodic signal is related to current sleep propensity and accumulated sleep need. It is unclear from the present results whether aperiodic measures might predict subjective sleepiness, subjective (versus objective) ratings of sleep propensity or circadian-related measures of sleepiness. Similarly, it is unclear whether aperiodic measures might be predictive of fatigue or time on task effects. These are all useful endeavours for future research.

Regarding the sleep EEG, SO-spindle coupling during NREM sleep differed significantly between symptomatic (MSLT+, MSLT–) and non-symptomatic (Control) individuals, with coupling strength being higher for the non-symptomatic group. These differences offer potential mechanistic insights within the context of hypersomnolence and underlying sleep neurophysiology. Sleep spindles and their coupling to slow oscillations are often disrupted in clinical relative to healthy individuals (for review, see ^47^). For example, the duration, amplitude, and density of sleep spindles differ between those with and without Alzheimer’s disease (AD), while SO-spindle coupling strength is negatively associated with two hallmark features of AD, namely pathological tau and *β*-amyloid aggregates ^63^. Poor or disrupted sleep is predictive of tau and *β*-amyloid (Aβ) accumulation, particularly within medial frontal regions and the hippocampal complex ^33^. From this perspective, individuals with EDS, who often report disrupted sleep, may be at greater risk of developing AD, a prediction supported by a recent study of 283 older individuals with EDS ^64^. Here, it was shown that baseline EDS predicted increased longitudinal Aβ accumulation. Given that our sample of clinically presenting patients were aged between 18 and 78 years (M_Age_ = 37.25 years), SO-spindle coupling strength may not only serve as a diagnostic tool in hypersomnolence treatment, but also a potential early marker of Aβ accumulation. Further, as SO-spindle coupling is plastic, if SO-spindle coupling is found to be casually related to EDS, brain stimulation (e.g., transcranial direct current stimulation) may serve as an alternative treatment method for individuals with hypersomnolence.

The development and validation of alternative methods for the diagnosis and assessment of hypersomnolence is an important endeavour for sleep research and medicine. The MSLT, currently the gold standard diagnostic procedure, is resource heavy necessitating a formal sleep laboratory as well as a full night and day of patient and laboratory scientist time. In contrast, the aperiodic slope may be a quick and efficient marker of sleepiness that is easier to obtain in a clinic environment thus enabling faster evaluation of EDS. Such a diagnostic tool could also play a role in serial monitoring over time, something that the MSLT struggles with given issues with cost, and test re-test reliability. Further research is required to explore these potential applications. Finally, it is also critical to highlight that the measurement of sleepiness via the MSLT and aperiodic components of the EEG as reported here does not relate to other measures of sleep and wake drive, such as vigilance, subjective sleepiness, or fatigue, which are typically measured with the maintenance of wakefulness test (MWT; ^65^). From this perspective, examining the aperiodic component of the EEG as it relates to the propensity to fall asleep versus the ability to maintain wakefulness may reveal similar or dissociable mechanisms of sleep and wake in the human brain.

The significance of sleepiness has recently been highlighted in experimental works which have linked the phenomenon to the E/I balance in neuronal functioning. From this, we have used a biological measure of E/I balance (the aperiodic slope) to classify EDS patients from healthy controls. We have also demonstrated potentially functionally relevant differences between these groups, in our observation of differences in SO-spindle coupling strength. This invites further fields of inquiry in terms of basic sleep research and medicine, which may lead us to a better understanding of sleep and sleepiness in health and disease.

## Acknowledgements

We thank Michelle Bay and Adelene Chin for assistance with PSG and MSLT data extraction. Thank you also to the patients from the Royal Adelaide Hospital and participants from the Cognitive Neuroscience Laboratory at the University of South Australia.

## Author Contribution

Conceptualization by all authors. Data curation by all authors. Data pre-processing and analysis by Z.C. Writing – original draft by Z.C. and A.C.; writing – review & editing by all authors; data visualization by Z.C.

## Financial Disclosure

None.

## Non-Financial Disclosure

None.

## Notes

### Competing Interest Statement

The authors have declared no competing interest.

