## supplementary material for "Resting-state aperiodic neural activity as a novel objective marker of daytime somnolence"

**S1.** Summary of linear mixed-effects regression examining aperiodic slope

|  | **Aperiodic Slope** | | |
| --- | --- | --- | --- |
| *Predictors* | *Estimates* | *CI* | *p* |
| (Intercept) | -2.38 | -2.51 – -2.24 | **<0.001** |
| Condition [Clinical] | 0.07 | -0.02 – 0.15 | 0.128 |
| Condition [Healthy] | 0.19 | 0.10 – 0.27 | **<0.001** |
| Sex [Female] | -0.04 | -0.12 – 0.04 | 0.332 |
| Age | 0.00 | 0.00 – 0.01 | **<0.001** |
| **Random Effects** | | | |
| σ^2^ | 0.02 | | |
| τ_00_ _subj_ | 0.02 | | |
| τ_00_ _Channel_ | 0.01 | | |
| ICC | 0.62 | | |
| N _subj_ | 85 | | |
| N _Channel_ | 5 | | |
| Observations | 425 | | |
| Marginal R^2^ / Conditional R^2^ | 0.233 / 0.712 | | |

**S2.** Summary of linear mixed-effects regression examining aperiodic intercept

|  | **Aperiodic Intercept** | | |
| --- | --- | --- | --- |
| *Predictors* | *Estimates* | *CI* | *p* |
| (Intercept) | -21.18 | -22.10 – -20.27 | **<0.001** |
| Condition [Clinical] | -0.46 | -1.05 – 0.13 | 0.125 |
| Condition [Healthy] | -0.75 | -1.33 – -0.17 | **0.012** |
| Sex [Female] | 0.08 | -0.45 – 0.61 | 0.764 |
| Age | -0.03 | -0.04 – -0.02 | **<0.001** |
| **Random Effects** | | | |
| σ^2^ | 0.41 | | |
| τ_00_ _subj_ | 1.18 | | |
| τ_00_ _Channel_ | 0.35 | | |
| ICC | 0.79 | | |
| N _subj_ | 85 | | |
| N _Channel_ | 5 | | |
| Observations | 425 | | |
| Marginal R^2^ / Conditional R^2^ | 0.179 / 0.825 | | |

**S3.** Summary of regression examining ESS

|  | **ESS** | | |
| --- | --- | --- | --- |
| *Predictors* | *Estimates* | *CI* | *p* |
| (Intercept) | 9.26 | 5.11 – 13.41 | **<0.001** |
| Condition [Clinical] | 1.88 | -0.73 – 4.49 | 0.156 |
| Condition [Healthy] | -5.40 | -8.56 – -2.25 | **0.001** |
| Sex [Female] | 0.05 | -2.79 – 2.89 | 0.972 |
| Age | 0.03 | -0.04 – 0.09 | 0.418 |
| Observations | 76 | | |
| R^2^ / R^2^ adjusted | 0.249 / 0.207 | | |

**S4.** Summary of linear mixed-effects regression examining slow oscillation density

|  | **SO Density** | | |
| --- | --- | --- | --- |
| *Predictors* | *Estimates* | *CI* | *p* |
| (Intercept) | 17.48 | 12.52 – 22.45 | **<0.001** |
| Group [MSLT+] | -0.10 | -2.64 – 2.44 | 0.939 |
| Group [Control] | 1.40 | -1.66 – 4.46 | 0.369 |
| Age | -0.25 | -0.33 – -0.18 | **<0.001** |
| **Random Effects** | | | |
| σ^2^ | 8.74 | | |
| τ_00_ _subj_ | 21.19 | | |
| τ_00_ _Channel_ | 13.46 | | |
| ICC | 0.80 | | |
| N _Channel_ | 4 | | |
| N _subj_ | 75 | | |
| Observations | 295 | | |
| Marginal R^2^ / Conditional R^2^ | 0.301 / 0.859 | | |

**S5.** Summary of linear mixed-effects regression examining slow oscillation peak-to-peak amplitude

|  | **SO PTP** | | |
| --- | --- | --- | --- |
| *Predictors* | *Estimates* | *CI* | *p* |
| (Intercept) | 129.43 | 116.66 – 142.19 | **<0.001** |
| Group [MSLT+] | 2.77 | -5.12 – 10.67 | 0.490 |
| Group [Control] | 3.27 | -6.22 – 12.76 | 0.498 |
| Age | -0.55 | -0.79 – -0.31 | **<0.001** |
| **Random Effects** | | | |
| σ^2^ | 184.90 | | |
| τ_00_ _subj_ | 177.91 | | |
| τ_00_ _Channel_ | 52.97 | | |
| ICC | 0.56 | | |
| N _Channel_ | 4 | | |
| N _subj_ | 75 | | |
| Observations | 295 | | |
| Marginal R^2^ / Conditional R^2^ | 0.167 / 0.630 | | |

**S6.** Summary of linear mixed-effects regression examining slow oscillation slope

|  | **SO Slope** | | |
| --- | --- | --- | --- |
| *Predictors* | *Estimates* | *CI* | *p* |
| (Intercept) | 527.24 | 447.04 – 607.44 | **<0.001** |
| Group [MSLT+] | 15.28 | -22.27 – 52.84 | 0.424 |
| Group [Control] | 33.41 | -11.75 – 78.57 | 0.146 |
| Age | -3.53 | -4.68 – -2.38 | **<0.001** |
| **Random Effects** | | | |
| σ^2^ | 3336.36 | | |
| τ_00_ _subj_ | 4244.86 | | |
| τ_00_ _Channel_ | 4030.16 | | |
| ICC | 0.71 | | |
| N _Channel_ | 4 | | |
| N _subj_ | 75 | | |
| Observations | 295 | | |
| Marginal R^2^ / Conditional R^2^ | 0.245 / 0.783 | | |

**S7.** Summary of regression examining slow oscillation-spindle coupling density

|  | **Coupling Density** | | |
| --- | --- | --- | --- |
| *Predictors* | *Estimates* | *CI* | *p* |
| (Intercept) | -12.54 | -16.92 – -8.16 | **<0.001** |
| Group [[MSLT-]] | -0.25 | -1.31 – 0.81 | 0.640 |
| Group [[MSLT+]] | -0.20 | -1.39 – 1.00 | 0.746 |
| Age | -0.13 | -0.19 – -0.08 | **<0.001** |
| Stage | 9.80 | 8.26 – 11.34 | **<0.001** |
| Observations | 151 | | |
| R^2^ / R^2^ adjusted | 0.575 / 0.563 | | |

**S8.** Summary of regression examining slow oscillation-spindle coupling strength

|  | **Coupling Strength** | | |
| --- | --- | --- | --- |
| *Predictors* | *Estimates* | *CI* | *p* |
| (Intercept) | 0.20 | 0.19 – 0.21 | **<0.001** |
| Group[MSLT-] | -0.02 | -0.02 – -0.02 | **<0.001** |
| Group[MSLT+] | -0.02 | -0.02 – -0.02 | **<0.001** |
| Age | -0.00 | -0.00 – 0.00 | 0.377 |
| Sex | -0.00 | -0.01 – 0.00 | 0.210 |
| Stage | -0.00 | -0.00 – 0.00 | 0.301 |
| **Random Effects** | | | |
| σ^2^ | 0.00 | | |
| τ_00_ _subj_ | 0.00 | | |
| ICC | 0.57 | | |
| N _subj_ | 77 | | |
| Observations | 151 | | |
| Marginal R^2^ / Conditional R^2^ | 0.820 / 0.922 | | |

**S9.** Summary of regression examining slow oscillation-spindle coupling phase

|  | **Coupling Phase** | | |
| --- | --- | --- | --- |
| *Predictors* | *Estimates* | *CI* | *p* |
| (Intercept) | 0.70 | 0.32 – 1.08 | **<0.001** |
| Group[MSLT-] | 0.28 | 0.17 – 0.38 | **<0.001** |
| Group[MSLT+] | 0.14 | 0.02 – 0.26 | **0.020** |
| Age | -0.00 | -0.01 – 0.00 | 0.102 |
| Stage | -0.19 | -0.32 – -0.06 | **0.003** |
| **Random Effects** | | | |
| σ^2^ | 0.16 | | |
| τ_00_ _subj_ | 0.03 | | |
| ICC | 0.18 | | |
| N _subj_ | 77 | | |
| Observations | 150 | | |
| Marginal R^2^ / Conditional R^2^ | 0.278 / 0.408 | | |
